# Genomic analysis reveals major determinants of *cis*-regulatory variation in *Capsella grandiflora*

**DOI:** 10.1101/034025

**Authors:** Kim A. Steige, Benjamin Laenen, Johan Reimegård, Douglas Scofield, Tanja Slotte

**Affiliations:** Dept. of Ecology and Genetics, Evolutionary Biology Centre, Uppsala University, Norbyv. 18D, 75236 Uppsala, SWEDEN; Science for Life Laboratory, Dept. of Ecology, Environment and Plant Sciences, Stockholm University, Lilla Frescati, SE-10691 Stockholm, SWEDEN; Science for Life Laboratory, Dept. of Cell and Molecular Biology, Uppsala University, Box 596, 75124 Uppsala, SWEDEN; Uppsala Multidisciplinary Center for Advanced Computational Science, Department of Information Technology, Uppsala University, Box 137, Uppsala 751 05, SWEDEN

**Author notes:** These authors contributed equally to the work (Kim A. Steige, Benjamin Laenen).

**Keywords:** allele-specific expression, distribution of fitness effects, purifying selection, positive selection, transposable elements, gene-body methylation

## Abstract

Understanding the causes of *cis*-regulatory variation is a long-standing aim in evolutionary biology. Although *cis*-regulatory variation has long been considered important for adaptation, we still have a limited understanding of the selective importance and genomic determinants of standing *cis*-regulatory variation. To address these questions, we studied the prevalence, genomic determinants and selective forces shaping *cis*-regulatory variation in the outcrossing plant *Capsella grandiflora*. We first identified a set of 1,010 genes with common *cis*-regulatory variation using analyses of allele-specific expression (ASE). Population genomic analyses of whole-genome sequences from 32 individuals showed that genes with common *cis*-regulatory variation are 1) under weaker purifying selection and 2) undergo less frequent positive selection than other genes. We further identified genomic determinants of *cis*-regulatory variation. Gene-body methylation (gbM) was a major factor constraining *cis*-regulatory variation, whereas presence of nearby TEs and tissue specificity of expression increased the odds of ASE. Our results suggest that most common *cis*-regulatory variation in *C. grandiflora* is under weak purifying selection, and that gene-specific functional constraints are more important for the maintenance of *cis*-regulatory variation than genome-scale variation in the intensity of selection. Our results agree with previous findings that suggest TE silencing affects nearby gene expression, and provide novel evidence for a link between gbM and *cis*-regulatory constraint, possibly reflecting greater dosage-sensitivity of body-methylated genes. Given the extensive conservation of gene-body methylation in flowering plants, this suggests that gene-body methylation could be an important predictor of *cis*-regulatory variation in a wide range of plant species.

## Significance

Despite long-standing interest in the contribution of *cis*-regulatory changes to adaptation, we still have a limited understanding of the selective importance and genomic determinants of *cis*-regulatory variation in natural populations. Using a combination of analyses of allele-specific expression and population genomic analyses, we investigate the selective forces and genomic determinants of *cis*-regulatory variation in the outcrossing plant species *Capsella grandiflora*. We conclude that gene-specific functional constraints shape *cis*-regulatory variation and that genes with *cis*-regulatory variation are under relaxed purifying selection compared to other genes. Finally, we identify a novel link between gene-body methylation and the extent of *cis*-regulatory constraint in natural populations.

## Introduction

Understanding the causes of regulatory variation is of major importance for many areas of biology and medicine (1). Much interest has centered on *cis*-regulatory variation, which has long been thought to be particularly important for adaptation (2–5). Like other quantitative traits, *cis*-regulatory variation is expected to be shaped by the interplay of mutation, selection, and drift. However, the relative importance of these forces remains unclear in most species.

Recently, prospects for quantifying *cis*-regulatory variation have greatly improved, and as a result, ample heritable cis-regulatory variation has been identified in many species (reviewed in (6)). This is resulting in a growing consensus that a large amount of standing *cis*-regulatory variation is under weak purifying selection (7–9). Clarifying why the impact of purifying selection varies across the genome is therefore important to understand the maintenance of *cis*-regulatory variation.

Variation in the intensity of purifying selection across the genome can result from differences in selective constraint that are due to the specific functions of the genes involved. For example, according to the dosage balance hypothesis, genes that encode interacting proteins are expected to experience stronger constraint than other genes (10). In yeast, there is empirical evidence that purifying selection on expression noise constrains regulatory evolution of dosage-sensitive genes (11–13) and in plants, dosage-sensitivity affects the retention of duplicate genes following whole-genome duplication (14). However, many other genomic features, including expression level, tissue specificity and gene-body methylation (gbM), are also known to be associated with constraint (15–18) and could affect *cis*-regulatory variation.

Variation in purifying selection can also result from broad, genome-scale forces that affect genes mainly as a result of their genomic environment, and not due to their specific function. For instance, in the self-fertilizing species *Caenorhabditis elegans*, variation in the impact of background selection across the genome had a major effect on the distribution of *cis*-regulatory variation across the genome (8). If background selection is important, then one might generally expect levels of *cis*-regulatory variation to be associated with recombination rate and/or gene density (19). At present however, the relative importance of gene-level constraint vs. genome-scale evolutionary forces for the distribution of *cis*-regulatory variation remains unclear in most species.

In this study, we have investigated the selective importance and genomic correlates of common *cis*-regulatory variation in the outcrossing crucifer species *Capsella grandiflora*. This species is particularly well suited for studying differences in the impact of selection across the genome, as it has relatively low population structure (20) and a large, stable effective population size (21, 22). Indeed, selection on both protein-coding (23) and regulatory regions (18) is highly efficient in *C. grandiflora,* and high levels of polymorphism enhance the power to detect *cis*-regulatory variation and quantify selection. Genomic studies are facilitated by the close relationship between *C. grandiflora* and the selfing species *Capsella rubella*, for which a genome sequence is available (22).

Here, we identified genes with common *cis*-regulatory variation in *C. grandiflora* based on analyses of allele-specific expression (ASE) in deep transcriptome sequencing data. To quantify the impact of positive and purifying selection on genes with *cis*-regulatory variation, we conducted population genomic analyses of high-coverage whole genome resequencing data from 32 *C. grandiflora* individuals. Finally, we identified genomic predictors of *cis*-regulatory variation. Our results show that there is pervasive *cis*-regulatory variation in *C. grandiflora*, and genes that harbor *cis*-regulatory variation are under weaker purifying selection and undergo less frequent positive selection than other genes. We find no evidence for a role of recombination rate or gene density in shaping *cis*-regulatory variation, suggesting that gene-specific variation in functional constraint is more important in this species. We further identify gbM as a major factor constraining *cis*-regulatory variation, whereas presence of nearby TEs and tissue specificity of expression increase the odds of ASE. Our results provide novel evidence for a link between gbM and *cis*-regulatory constraint, possibly reflecting greater dosage-sensitivity of body-methylated genes.

## Results

### Widespread cis-regulatory variation in C. grandiflora

To identify genes with *cis*-regulatory variation, we quantified allele-specific expression based on deep whole transcriptome sequencing data (total 93.5 Gbp with Q≥30) from flower buds and leaves of three *C. grandiflora* F1s (Supplementary Table S1). Each F1 harbored an average of about 235,700 high-confidence heterozygous coding SNPs, which were phased prior to analyses of ASE. After filtering, on average approximately 13,400 genes per F1 were amenable to ASE analyses (Table 1).

**Table 1.** Genes amenable to analysis of ASE in flower buds and leaves, and ASE results.

| Sample type | F1 | Analyzed<sup>1</sup> | ASE genes<sup>2</sup> | ASE prop.<sup>3</sup> | FDR<sup>4</sup> |
| --- | --- | --- | --- | --- | --- |
| Flower buds | 6.3 | 13521 | 3065 | 0.33 | 0.00134 |
|  | 7.2 | 14390 | 3829 | 0.36 | 0.00240 |
|  | 8.2 | 14232 | 3601 | 0.35 | 0.00198 |
| Leaves | 6.3 | 12390 | 3425 | 0.34 | 0.00182 |
|  | 7.2 | 13074 | 3749 | 0.39 | 0.00242 |
|  | 8.2 | 12796 | 3550 | 0.34 | 0.00195 |

<sup>1</sup>Genes with expression data for at least one replicate, and with a phased fragment containing at least three transcribed SNPs after filtering.
<sup>2</sup>Genes with posterior probability of ASE $\geq 0.95$ .
<sup>3</sup>Estimated proportion of genes with ASE
<sup>4</sup>False Discovery Rate

We assessed ASE using a Bayesian method (24), accounting for technical variation in allelic counts using high-coverage whole genome resequencing data for each F1 (mean coverage of 40x, total 26.6 Gbp with Q≥30; Supplementary Table S2). We estimated that a mean of 35% (range 33-39%) of analyzed genes show ASE in individual *C. grandiflora* F1s (Table 1). Similar proportions of genes had ASE in both leaves and flower buds (Table 1) and allelic expression biases were moderate for most genes with ASE, with strong allelic expression biases (0.2 ≤ ASE ratio ≥ 0.8) shown by an average of 5.1% of genes (Figure 1, Supplementary Figures S1 and S2).

**Figure 1.**
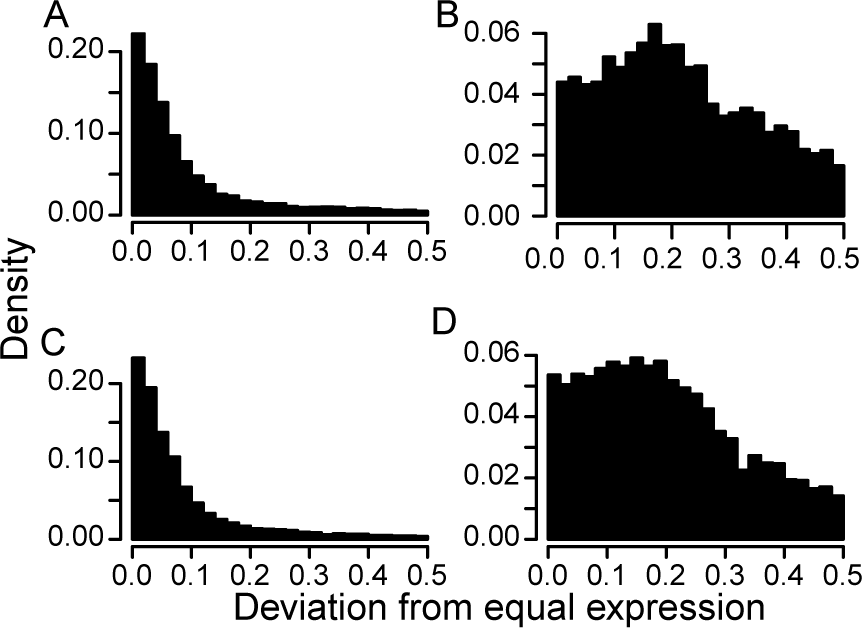
The extent of ASE in leaves (A, B) and flower buds (C, D) of one representative *C. grandiflora* F1. Panels A and C show the deviation from equal expression for all assayed genes. Genes with strong evidence for ASE (posterior probability of ASE ≥ 0.95) show stronger deviations from equal expression (B and D).

Out of a total of 11,532 genes that were amenable to analysis of ASE in all F1s, there were 1,010 genes that showed ASE in either leaves or flower buds, 313 genes showed ASE in flower buds but not leaves, 404 genes showed ASE in leaf samples but not flower buds, and 293 genes had ASE in both flower buds and leaves of all F1s (Supplementary Figure S3). Among the 1,010 genes with ASE leaves or flower buds of all F1s, one GO category, GO:0006952, “defense response” was significantly enriched at FDR ≤ 0.01. This was likely driven by genes with ASE in leaves, as there was no significant enrichment of Gene Ontology (GO) terms among genes with ASE in flower buds, whereas six biological process GO terms associated with photosynthesis and defense responses were significantly enriched (FDR ≤0.01) among genes with ASE in leaves (Supplementary Table S3). Among control genes, there was a nominally significant enrichment of genes in only two GO terms, protein binding (GO:0005515) and zinc ion binding (GO:0008270) (Weighted Fisher *P* ≤ 0.01), but this was not significant at FDR ≤ 0.01.

### Lower intensity of purifying selection on genes with cis-regulatory variation

To assess the impact of selection on genes showing *cis*-regulatory variation in *C. grandiflora*, we sequenced the genomes of 21 individuals from one population in the Zagory region of Greece (the ‘population sample’) as well as 12 individuals from separate populations across the species range (the ‘range-wide sample’) using 233.2 Gbp of high-quality (Q≥30) paired-end 100 bp Illumina reads and a mean coverage of 25x per individual (Supplementary Table S2). We called variants using GATK best practices and filtered genomic regions as previously described (25) to identify a total of 6,492,075 high-quality SNPs, most of which (5,240,485) were also segregating in the population sample.

We compared levels of polymorphism at genes that show ASE in all of our F1s (1,010 genes; ‘ASE genes’), using as a control set the 10,552 genes that were amenable to ASE analyses in all F1s but did not show significant ASE (termed ‘control genes’) (Supplementary Figure S3). To reduce bias resulting from the requirement of expressed polymorphisms for analyses of ASE, all population genetic analyses were conducted only on these paired gene sets, and genes that were not amenable to analysis of ASE were not included. ASE genes had elevated polymorphism levels compared to the control at all investigated site classes, as well as an elevated ratio of nonsynonymous to synonymous polymorphism (Table 2; Supplementary Table S4), suggesting that the impact of purifying selection might differ between ASE and control gene sets (Table 2; Supplementary Table S4).

**Table 2.** Population genetic summary statistics and divergence estimates for the different site classes, separately for ASE and control genes.

| Site class<sup>1</sup> | Gene set | Mean $\pi$ | Mean $\theta_W$ | $\pi_{\text{siteclass}}/\pi_{\text{4-fold}}$<sup>2</sup> | $d$ |
| --- | --- | --- | --- | --- | --- |
| 4-fold | ASE | 0.029 | 0.029 | NA | 0.16 |
|  | control | 0.023 | 0.024 | NA | 0.15 |
| 0-fold | ASE | 0.009 | 0.011 | 0.32 | 0.04 |
|  | control | 0.005 | 0.007 | 0.23 | 0.03 |
| 3'UTR | ASE | 0.018 | 0.021 | 0.62 | 0.13 |
|  | control | 0.014 | 0.018 | 0.62 | 0.12 |
| 5'UTR | ASE | 0.016 | 0.016 | 0.55 | 0.12 |
|  | control | 0.012 | 0.012 | 0.54 | 0.12 |
| 500 bp upstream | ASE | 0.017 | 0.020 | 0.6 | 0.16 |
|  | control | 0.015 | 0.019 | 0.68 | 0.15 |
| intron | ASE | 0.020 | 0.022 | 0.69 | 0.15 |
|  | control | 0.018 | 0.020 | 0.79 | 0.14 |

<sup>1</sup>Class of sites investigated, including 4-fold degenerate sites (4-fold), 0-fold degenerate sites (0-fold), 5'UTRs, 3'UTRs, 500 bp upstream of the TSS (500 bp upstream) and introns.
<sup>2</sup> Ratio of nucleotide diversity at focal site class to nucleotide diversity at 4-fold synonymous sites.

To quantify the impact of purifying selection on ASE genes and control genes, we used the DFE-alpha method (26, 27), which allows estimation of a gamma-distribution of negative fitness effects based on site frequency spectra (SFS) at putatively neutral and selected sites. We found that ASE genes have a significantly higher proportion of nearly neutral nonsynonymous mutations than control genes, as well as a significantly reduced proportion of nonsynonymous mutations under strong purifying selection (strength of purifying selection *N_e_s*>10) (Figure 2). This result applies broadly, both for the population and the range-wide samples, and when assuming a constant population size as well as after correcting for population size change (Supplementary Figure S4). The result also holds after controlling for differences in the expression level among genes with and without ASE (Supplementary Figure S5), and when classifying genes based on a single F1 individual (Supplementary Figure S6), suggesting that the results hold broadly for common *cis*-regulatory variation. Our results further remain unchanged after removing defense-response genes (GO:0006952) with ASE (Supplementary Figure S7) prior to DFE-alpha analyses, and thus strong balancing selection on these genes does not drive the patterns we observe.

**Figure 2.**
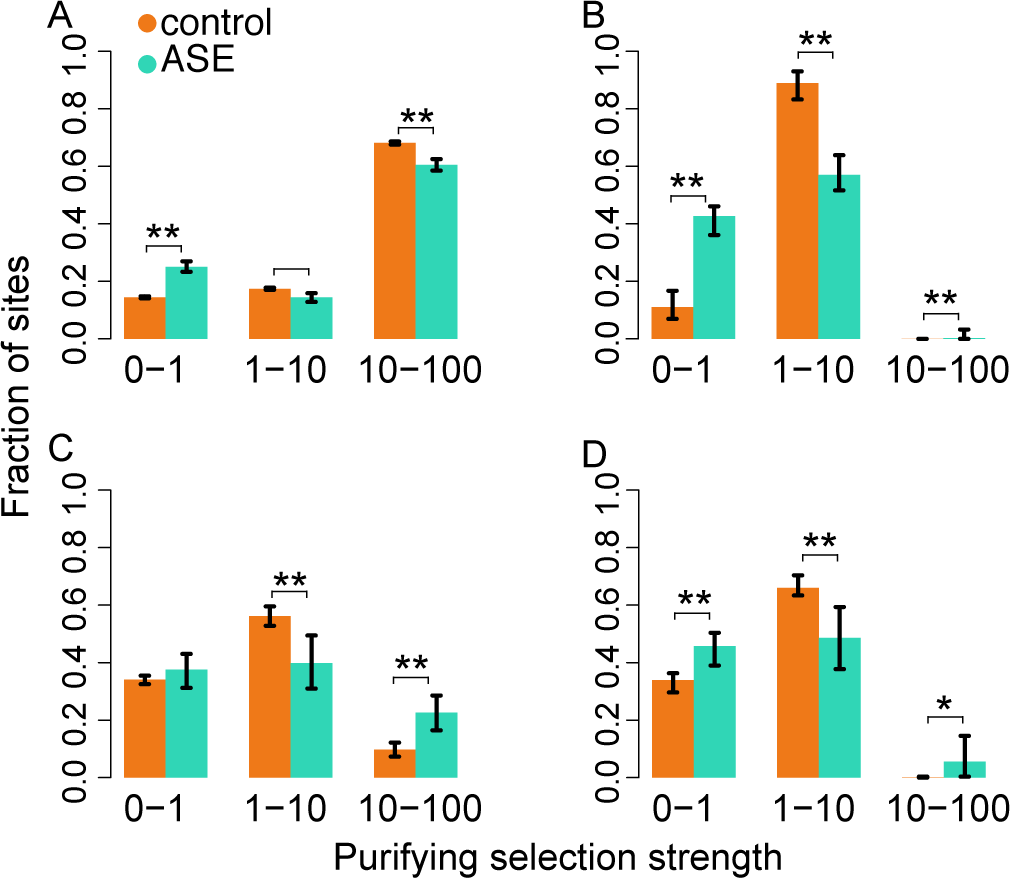
The impact of purifying selection differs between genes with and without ASE in *C. grandiflora.* The estimated proportion of mutations in each *N_e_s* bin of the distribution of negative fitness effects (DFE) is shown, with whiskers corresponding to 95% confidence intervals. The panels show the DFE for nonsynonymous sites (0-fold degenerate sites) (A), for introns (B), for promoter regions 500 bp upstream of the transcription start site (C) and for 3’-UTRs (D). Significance levels for comparisons of ASE and control genes, that were also amenable to analysis of ASE, are indicated by asterisks (*: P-value ≤ 0.05, ** P-value ≤ 0.01). These results are based on the population sample and the one-epoch model.

In contrast to the clear evidence for weaker purifying selection at nonsynonymous sites for genes with ASE, there were no significant differences in the DFE depending on ASE status at 5’-UTRs (Supplementary Figure S8). For introns, results were inconsistent, with some but not all analyses pointing to weaker purifying selection on control genes (Figure 2, Supplementary Figure S9, Supplementary Table S5). This could suggest that patterns of selection differ among coding and noncoding regions. However, at other noncoding regions than introns, such as promoter regions 500 bp upstream of the TSS and at 3’-UTRs, there was some evidence for relaxed purifying selection at ASE genes (Figure 2; Supplementary Figures S10-S11, Supplementary Table S5). These results held only under the 1-epoch model, which could in part be due to a lack of power, as regulatory motifs are expected to make up a small fraction of the analyzed sites. Consistent with this, we infer weaker purifying selection on upstream regions and UTRs than on nonsynonymous mutations (Supplementary Figures S8-S11; Supplementary Table S5).

### Genes with cis-regulatory variation undergo less frequent adaptive evolution

To investigate the impact of positive selection on genes with and without ASE we obtained estimates of *ω_a_*, the rate of adaptive substitutions relative to neutral divergence (28) in DFE-alpha. For this purpose, we relied on genome-wide divergence between *Capsella* and *Arabidopsis*, with 4-fold synonymous sites considered to be evolving mainly neutrally (see Methods for details). Using this method, we find that ASE genes show a significantly lower proportion of adaptive nonsynonymous substitutions than control genes (Figure 3). In contrast, we found no significant differences in *ω_a_* among ASE genes or control genes for UTRs or regions 500 bp upstream of the TSS (Supplementary Table S5). Second, we estimated *α*, the proportion of adaptive fixations in the selected site class, based on the approximate method of (29), designed to yield accurate estimates in the presence of linked selection. Results generated with this method were consistent with DFE-alpha, with a significantly lower estimate of the proportion of adaptive nonsynonymous substitutions at genes with *cis*-regulatory variation than at control genes in *C. grandiflora* (Figure 3).

**Figure 3.**
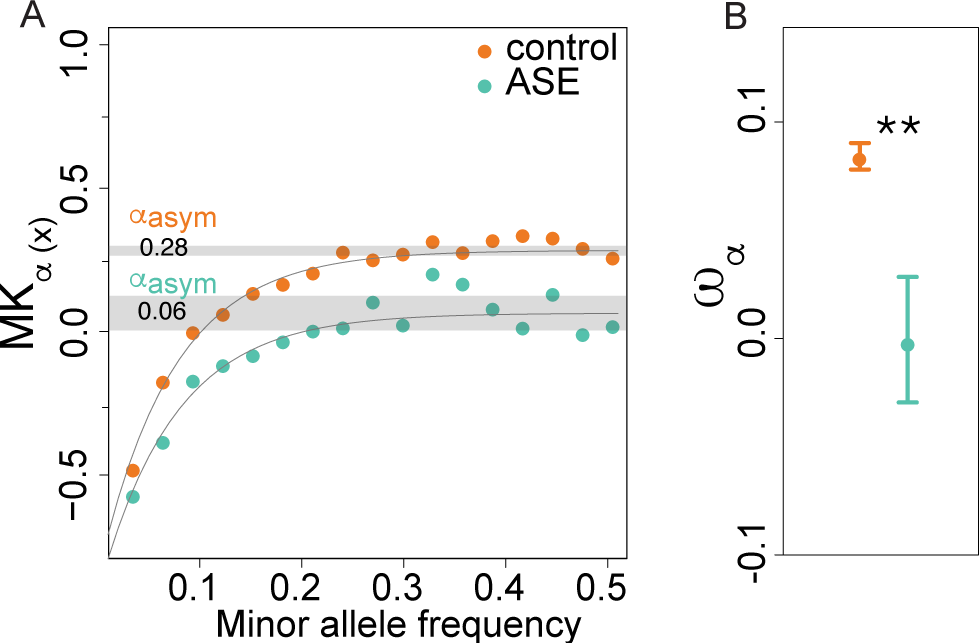
A lower proportion of adaptive nonsynonymous fixations at genes with ASE. (A) Estimation of α using the asymptotic method of Messer and Petrov (2013), which fits an exponential function to estimates of α based on polymorphisms at different frequencies. Orange dots show values for control genes, and green dots show values for genes with ASE. The grey shaded area indicates 95% confidence intervals. The point estimate for genes with and without ASE is 0.06 vs 0.28, respectively. (B) The estimated proportion of adaptive fixations relative to 4-fold synonymous substitutions (*ω*_a_) for genes with and without ASE. Whiskers correspond to 95% confidence 614 intervals, and significance levels for comparisons of ASE and control genes are indicated by asterisks (*: P-value ≤ 0.05, ** P-value ≤ 0.01).

### Determinants of cis-regulatory variation in C. grandiflora

To identify genomic factors and potential drivers of *cis*-regulatory variation, we conducted logistic regression analyses with presence/absence of ASE as the response variable. We included a total of 12 predictor variables, chosen to include proxies for variation in mutation rate, recombination rate, gene density, expression level and degree of constraint, which could be expected to affect levels of *cis*-regulatory variation (see Methods for details). The best-fit model retained eight of these predictor variables (Table 3). In this model, gbM had the greatest effect on *cis*-regulatory variation, resulting in a reduction of 49% in the odds of observing ASE (Table 3), whereas the presence of polymorphic TEs within 1 kb of the gene also had a substantial effect, increasing the odds of ASE by 38%, followed in turn by tissue specificity of expression, promoter diversity, expression level, gene length and nonsynonymous/synonymous polymorphism, all of which increased the odds of ASE (Table 3). Including network connectivity improved model fit, although the effect was not individually significant (Table 3). Notably, gene density and recombination rate, which affect the intensity of linked selection, were not included in the best-fit model. Similar results were obtained in an analysis that followed the approach of (30) to ensure orthogonality of predictors by using principal components of all continuous predictors in logistic regression analyses (Supplementary Tables S6 and S7). These analyses suggest that variation in gene-specific constraint are important for shaping the distribution of *cis*-regulatory variation across the *C. grandiflora* genome, and that gbM and presence of nearby TEs are strong predictors of *cis*-regulatory constraint.

**Table 3.** The best-fit logistic regression model (AIC=3086.9) predicting ASE from genomic features. Regression coefficients and their standard error, z-statistics and associated P-values, and odds ratios (OR) are shown. Coexpression module size and gene length were included in the best-fit model, but did not have individually significant effects.

| Model parameter | Coeff. (SE) | z value | P-value | OR |
| --- | --- | --- | --- | --- |
| Gene-body methylation | -0.67 (0.20) | -3.41 | <10<sup>-3</sup> | 0.51 |
| $\pi_N/\pi_S$ | 0.08 (0.04) | 2.25 | 0.024 | 1.09 |
| Expression level | 0.20 (0.06) | 3.31 | <0.001 | 1.22 |
| Promoter polymorphism | 0.21 (0.05) | 4.45 | <10<sup>-3</sup> | 1.23 |
| Tissue specificity | 0.30 (0.06) | 5.03 | <10<sup>-3</sup> | 1.35 |
| TE within 1 kb | 0.32 (0.13) | 2.50 | 0.013 | 1.38 |
| Coexpression module size | -0.08 (0.05) | -1.59 | NS | 0.92 |
| Gene length | 0.08 (0.06) | 1.49 | NS | 1.09 |
| Intercept | -2.60 (0.06) | -42.91 | <10<sup>-3</sup> | 0.07 |

## Discussion

Our results show that genes that harbor common *cis*-regulatory variation in *C. grandiflora* are under weaker purifying selection, and experience less frequent positive selection than other genes. We further find that gene-specific features that are likely to reflect the degree of functional constraint and mutational input are better predictors of *cis*-regulatory variation than those that are expected to shape the broad impact of linked selection across the genome. These functional constraints do not appear to limit the potential for adaptation at coding sequences, as positive selection had a greater impact on coding divergence at genes that did not exhibit common *cis*-regulatory variation in *C. grandiflora*.

Our findings support the view that most standing *cis*-regulatory variation in natural populations is weakly deleterious (7), and our robust inference of relaxed purifying selection on genes with common *cis*-regulatory variation agrees well with those of a recent eQTL mapping study in *C. grandiflora* (9). Our inference of relaxed purifying selection on genes with common *cis*-regulatory variation do not appear to be driven by balancing selection or conditional neutrality affecting a subset of defense-related genes that show ASE, as our results remain unchanged after removing such genes.

The major association between gbM and *cis*-regulatory constraint that we detected is particularly interesting, because the function of gbM is currently unclear (31, 32). The conservation of gbM of orthologs in very distantly related plant species suggests that gbM has functional importance, but intriguingly, some plants lack gbM (31–33). Body-methylated genes tend to be longer than other genes, expressed at intermediate levels, evolve slowly at the sequence level (17, 34, 35), and are stably expressed under different conditions (36). A recent study found that *A. thaliana* from northern Sweden show elevated gene body methylation, mainly due to *trans*-acting loci (36), but as far as we are aware, no study has directly linked gbM to *cis*-regulatory variation in natural plant populations.

It is possible that the associations we detected between genomic features and *cis*-regulatory variation are caused by underlying drivers that were not directly measured. One natural candidate is gene essentiality. However, while gbM is significantly associated with predicted gene essentiality (37) (Fisher exact test P<0.001), our results do not appear to be driven by essentiality, which was not retained in our best-fit logistic regression model for *cis*-regulatory variation. Instead, we hypothesize that selection for increased stability of expression of dosage-sensitive genes could underlie several of the associations we observe. Dosage-sensitive genes exhibit less expression noise (12, 38), show less variation in expression among tissues, and are expected to be part of larger regulatory network modules(10, 12). In our study, reduced tissue-specificity of expression and increased network connectivity were associated with a reduced likelihood of ASE. Furthermore, expression variation among three biological replicates of a *C. rubella* genotype (25) that likely represents mainly noise, is significantly lower for genes with no ASE than for those with ASE (median CV of FPKM=0.28 for genes with ASE, 0.18 for control genes, Wilcoxon rank-sum test, P-value<10^-5^). Finally, defense-related genes, which are thought to be dosage-insensitive in plants (39), were significantly enriched among genes with *cis*-regulatory variation in our study, whereas protein-binding genes were nominally enriched among control genes without ASE. Both promoter polymorphism and TE insertions, which can impact expression in several ways (40), might be expected to be more likely to be tolerated near dosage-insensitive genes. Our results are therefore consistent with dosage sensitivity causing strong constraint on *cis*-regulatory variation and shaping the impact of positive and purifying selection on coding variation. Thus, similar functional constraints that shape duplicate gene retention after whole genome duplication (14) may also be key for the genomic distribution of *cis*-regulatory variation in natural plant populations. Future studies should explore the connection between dosage-sensitivity, gbM, and *cis*-regulatory variation in greater detail across a wider range of plant species.

## Materials and Methods

### Plant Material

For analyses of ASE, we generated three intraspecific *C. grandiflora* F1s by crossing six individuals sampled across the range of *C. grandiflora* (Supplementary Table S8). For population genomic analyses of *C. grandiflora*, we grew a single offspring from field-collected seeds of each of 32 plants (‘the population genomic sample’; Supplementary Table S9), representing 21 plants from one population from Greece (the ‘population sample’), and 11 additional plants from 11 separate Greek populations covering the species’ range. Together with an individual from the population sample, these represent a 12-plant ‘range-wide sample’. We grew plants at standard long-day conditions and collected leaf and mixed stage flower bud samples for RNA sequencing, and leaf samples for whole genome sequencing as previously described (25).

### Sample preparation and sequencing

We extracted total RNA from the intraspecific F1s using a Qiagen RNEasy Plant Mini Kit (Qiagen, Hilden, Germany). RNAseq libraries were constructed using the TruSeq RNA v2 kit. For genomic resequencing, we extracted genomic DNA using a modified CTAB extraction method. Whole genome sequencing libraries with an insert size of 300-400 bp were prepared using the TruSeq DNA v2 protocol. Sequencing of 100bp paired-end reads was performed on an Illumina HiSeq 2000 instrument (Illumina, San Diego, CA, USA). All sequence data has been submitted to the European Bioinformatics Institute (www.ebi.ac.uk), with study accession numbers: PRJEB12070 and PRJEB12072.

### Sequence quality and trimming

RNA and DNA reads from the F1s were trimmed as previously described (25). Adapters and low quality sequence were trimmed using CutAdapt 1.3. We analyzed genome coverage using BEDTools v.2.17.0 (41) and removed potential PCR duplicates using Picard v.1.92 (http://picard.sourceforge.net).

### Read mapping, variant calling and filtering

We mapped RNAseq reads from the F1s to the v1.0 reference *C. rubella* assembly (22) using STAR v.2.3.0.1 (42) with default parameters. For genomic reads from F1s, we mapped reads with STAR as in (25). Genomic reads from the population genomic sample were mapped using BWA-MEM v.0.7.12 (43) using default parameters and the –M flag.

Variant calling was done using GATK v. 2.5-2 UnifiedGenotyper (44) according to GATK best practices (45, 46). We conducted duplicate marking, local realignment around indels and recalibrated base quality scores using a set of 1,538,085 SNPs identified in *C. grandiflora* (18) as known variants, and retained only SNPs considered high quality by GATK.

We removed centromeric and pericentromeric regions where we have low confidence in our variant calls, and prior to ASE analysis, we conducted additional filtering of SNPs as in (25). Using this procedure, we identified an average of 235,719 heterozygous coding SNPs in 17,973 genes in each F1. For population genomic analyses, we further filtered all genomic regions annotated as repeats using RepeatMasker 4.0.1, and removed sites with extreme coverage (DP < 15 or DP > 200) and too many missing individuals (≥20%) using VCFtools (47). Indels and non-biallelic SNP were also pruned prior to analysis.

### Phasing

Prior to ASE analysis, we conducted read-backed phasing of genomic variants in F1s using GATK v. 2.5-2 ReadBackPhasing (-phaseQualityThresh 10). RNAseq data from all F1s were subsequently phased by reference to the phased genomic variants. Read counts for all phased fragments were obtained using Samtools mpileup. This resulted in a mean number of 31,313 contiguous phased fragments per F1 (Table 1).

To validate our phasing procedure, we compared the phased fragments, based on reads, with the phased chromosomes, based on heritage, in three interspecific *C. grandiflora* x *C. rubella* F1s from (25). For most genes, over 95% of SNPs were correctly phased in the interspecific F1s, demonstrating that our phasing procedure is reliable (Supplementary Figure S13; Supplementary Figure S14).

### Analyses of allele-specific expression

We analyzed ASE using a hierarchical Bayesian method which requires phased data, in the form of read counts at heterozygous SNPs for both genomic and transcriptomic data (24). Genomic read counts are used to obtain an empirical estimate of technical variation which is then used in analyses of the RNAseq data. We used this method to obtain estimates of the posterior probability and degree of ASE, for the longest phased fragment per gene with at least three transcribed SNPs. We analyzed ~14,000 reliably expressed genes for ASE in flower buds, and ~13,400 genes in leaves (Table 1). All analyses were run in triplicate, and we checked MCMC convergence by comparing parameter estimates from independent runs with different starting points, and by assessing mixing. Runs were completed using the pqR version of R (http://www.pqr-project.org) for 200,000 generations or a maximum runtime of 10 days, with the first 10% of each run discarded as burn-in.

### Population genomic analyses

To assess whether patterns of polymorphism differ among ASE and control genes, we tested for a difference in median levels of polymorphism and Tajima’s *D* in the *C. grandiflora* population sample, using Mann-Whitney U-tests, with Benjamini-Hochberg correction for multiple comparisons. Estimates of nucleotide diversity (*π*), Watterson’s theta (*θ_W_*) and Tajima’s *D* (*D*_T_) were obtained using custom R scripts by BL. Separate estimates were obtained for 6 classes of sites: 4-fold degenerate sites, 0-fold degenerate sites, 3’- and 5’-untranslated regions (UTRs), introns, and intergenic regions 500 bp upstream of the transcription start site (TSS).

### Selection on genes with ASE

To test whether there was evidence for a difference in the strength and direction of natural selection on ASE and control genes, we first estimated the distribution of fitness effects (DFE) as in (26), and the proportion of adaptive substitutions relative to the total number of synonymous substitutions (*ω_a_*) using the method of (28). The DFE was estimated under a constant population size model and under a model with stepwise population size change. We obtained confidence intervals for our estimates of three bins of the DFE (0<*N_e_s*<1; 1<*N_e_s*<10; 10<*N_e_s*) and for *α* and *ω_a_* by resampling genes in 200 bootstrap replicates and tested for a difference in the DFE, and *ω_a_* among sets of genes with ASE and control genes, as in (27). Separate estimates were obtained for 0-fold degenerate sites, 3’- and 5’-untranslated regions (UTRs), introns, and promoter regions 500 bp upstream of the TSS likely enriched for regulatory elements, using 4-fold degenerate sites as neutral standard. For estimates of *α* and *ω_a_*, we relied on divergence to *Arabidopsis*; specifically, we generated a whole genome alignment using lastz v. 1.03.54, with chaining of *C. rubella*, *A. thaliana* and *A. lyrata* as described in (48), and counted divergence differences and sites as in (18). DFE-alpha analyses were run using Method I (27).

To assess the effect of expression level on our DFE-alpha inference, we selected genes among the control set of genes to match the distribution of expression level of ASE gene by resampling the control genes to match the distribution of expression levels in the ASE gene set. Purifying selection and positive selection were then re-evaluated in DFE-alpha, using the resampled control gene set and the ASE set. To assess whether our results were robust to the sampling strategy for ASE analyses, we based our classification of ASE and control genes based on a single F1 individual and repeated the DFE analyses. To investigate whether our results could be driven by the inclusion of defense-related genes, we removed genes annotated as defense-response genes (GO:0006952), and repeated the DFE-alpha analyses.

### Genomic determinants of cis-regulatory variation

We assessed the relative importance of a number of genomic features for presence/absence of ASE using logistic regression, on a set of genes that was restricted to those for which we could assess ASE. We included the following genomic features that may affect linked selection: recombination rate and gene density (in 50 kb windows). Gene density were based on the annotation of *C. rubella* v1.0 reference genome (22). We obtained recombination rates per 50kb windows based on 878 markers from (49) by fitting a smooth spline. We further included gene length, tissue specificity (*t*;(16)), expression level (log FPKM values), and as a proxy for mutation rate variation, we included 4-fold synonymous divergence to Arabidopsis (*d_S_*). Because promoter polymorphism may cause *cis*-regulatory variation, we included nucleotide diversity (*π*) for the region 500bp upstream of the TSS. We included nonsynonymous/synonymous nucleotide diversity (*π_N_*/*π_S_*) to reflect the level of constraint at the coding sequence level. According to the dosage balance hypothesis, genes in smaller co-expression modules may be under reduced regulatory constraint. We therefore included information on *A. thaliana* co-expression module size (37) in our analyses. We further included information on the presence of retained paralogs from the Brassicaceae α whole genome duplication or the β and γ whole genome duplication (37). We identified a set of genes with gbM in both *C. rubella* (33) and *A. thaliana* (17), which are highly likely to also harbor gbM in *C. grandiflora*. Finally, we included information on polymorphic TEs within 1 kb of genes in the range-wide sample. We identified TE insertions in our range-wide sample as in (25), except that we required a minimum of 5 reads to call a TE insertion. All continuous variables were centered and scaled prior to logistic regression, with model selection using a stepwise AIC procedure with backward and forward selection of variables to find the best-fit model (Table 3). We repeated the analysis using an analysis strategy which is superior to partial correlation analysis and robust in the presence of noisy genomic data and multicollinearity of predictor variables (30). We used a set of orthogonal predictor variables obtained by identifying principal components for a data set including all the continuous variables using the “pls” package in R, well as gene body methylation and presence of heterozygous TEs as binary factors, and conducted model selection as described above.

## Acknowledgements

We thank Lauren McIntyre and Stephen Wright for valuable discussion, Daniel Halligan for sharing scripts for DFE-alpha analyses and Daniel Skelly for advice on ASE analyses. We thank Veronika Scholz and Michael Nowak for bioinformatic assistance, and Julia Dankanich and Cindy Canton for assistance with experiments and lab work. Sequencing was performed by the SNP&SEQ Technology Platform in Uppsala. The computations were performed on resources provided by SNIC through Uppsala Multidisciplinary Center for Advanced Computational Science (UPPMAX) under projects b2012122 and b2012190. This study was funded by grants from the Swedish Research Council, the Nilsson-Ehle foundation, the Magnus Bergvall foundation, and the Erik Philip-Sörensen foundation to T.S.

